# Selecting among three basic fitness landscape models: additive, multiplicative and stickbreaking

**DOI:** 10.1101/150763

**Authors:** Craig R. Miller, James T. Van Leuven, Holly A. Wichman, Paul Joyce

## Abstract

Fitness landscapes map genotypes to organismal fitness. Their topography depends on how mutational effects interact–epistasis–and is important for understanding evolutionary processes such as speciation, the rate of adaptation, the advantage of recombination, and predictability versus stochasticity of evolution. The growing amount of empirical data has made it possible to better test landscape models empirically. We argue that this endeavor will benefit from the development and use of meaningful null models against which to compare more complex models. Here we develop statistical and computational methods for fitting fitness data from mutation combinatorial networks to three simple models: additive, multiplicative and stickbreaking. We employ a Bayesian framework for doing model selection. Using simulations, we demonstrate that our methods work and we explore their statistical performance: bias, error, and the power to discriminate among models. We then illustrate our approach and its flexibility by analyzing several previously published datasets. An R-package that implements our methods is available in the CRAN repository under the name *Stickbreaker.*

## 1. Introduction

The fitness landscape is a modeling framework that maps DNA or protein sequence variants to fitness ([1], [2], [3], [4]). Adjacent locations on a plane represent genomes that differ by one mutational event. The fitness of each genotype is envisioned as forming a surface above the plane. Fisher’s Geometric model is closely related. There, the plane represents phenotype space (rather than sequence space) and again the surface above is fitness ([5], [3], [6], [7], [8]). In reality, of course, the genotype (or phenotype) plane is often highly dimensional; a two dimensional plane with a fitness surface above is used mainly because it begets the landscape metaphor and makes the model easier to conceptualize.

Understanding the topography of the fitness landscape is important. It determines the extent to which recombination confers benefits, which bears on the potential advantages of sex ([9], [10], [11]); it has consequences for reproductive isolation as a mechanism for speciation ([12], [13], [14]); it dictates how stochastic vs predictable evolution is ([15], [16], [17], [18]); it plays a major role in how likely and at what speed adaptation is to find a highly optimal solution ([19], [20], [21]). But developing an understanding of real fitness landscapes is a serious challenge. First, the space is staggeringly vast and estimating its shape from a small sample of the space can be misleading ([22]). Even in a tiny viral genome of 5000 bases, there are 5000 x 3 = 15,000 possible first step DNA substitutions and the number of different genotypes with, say, just five mutations is on the order of 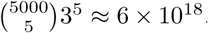. The number of unique pathways to each of these adds severals more orders of magnitude: 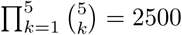. Second, fitness on the landscape in real populations is rarely fixed; it shifts over time due to biotic and abiotic changes in the environment ([23], [24], [25], [26]). Third, the underlying biology of this space is complex and this makes developing models based on the biology difficult.

In the face of these challenges, researchers have pursued two major strategies to studying fitness landscapes: theoretical and empirical. An extensive body of theory has been developed that is based on various assumptions about relevant features such as the number and distribution of mutational effects on fitness, how mutations interact (epistasis), and the mutation-selection dynamics at work in the population (e.g. [27], [2], [23], [28], [29], [30], [6], [31], [20], [32], [33] are good examples among a large body of literature). In the second approach, empirical data about the fitness landscape are collected ([4]). Given the vast scope of sequence space, these studies have necessarily focused data collection on small regions of the landscape. One way to obtained an especially detailed view of the landscape is to begin with a small set of mutations, construct all combinations of them and then measure their phenotypes and/or fitnesses (e.g. [34], [35], [36], [16], [37], [38], [39], [40], [41], [42], [43], [44], [8]). In the landscape metaphor, this maps out all possible mutational pathways between the wild type and the genotype with all mutations included. One common variation of this approach is to use pairs of mutations and engineer the two single mutants and double mutant genotypes (e.g. [45], [7], [46]); this amounts to creating many two-step, 4-genotype networks. As tools for genetic engineering improve, these experimental approaches are becoming increasingly feasible for greater numbers of mutations and larger mutational networks (e.g., [47], [48], [49]).

There is growing momentum in the field to bridge the theoretical and empirical ([4]). Much of mutational combination data has been fit or compared to models–either in the original work, in later analysis papers, or both. One common approach has been to assume a null model (usually the additive or multiplicative model) and characterize epistasis as deviations from this (e.g. [45], [38], [40], [46]). Another approach has been to characterize the extent of sign epistasis in the data (the case where mutations switch from being individually deleterious to being beneficial in combination, or vice versa) and, using a model of population dynamics, examine the probabilities of different pathways in the network (e.g. [34], [16], [50], [42], [44]). Other studies have fit the data to landscape models. Among explicitly fitted models, one group is based on mapping genotypes to fitness and includes the Rough Mt. Fuji model, the NK model, the uncorrelated model (also known as the House of Cards) as well as models more tailored to the biology of the study system (e.g. [35], [51], [41], [39], [52], [53], [54]). The other family of fitted models is based on Fisher’s geometric model where mutations are assumed to have additive effects on phenotype and phenotypes map to fitness (e.g. [36], [55], [7], [56], [26], [8]).

We believe that this endeavor of fitting data to landscape models can be strengthened by more carefully considering and developing null models. More specifically, it has been generally overlooked that there are actually several equally simple fitness landscape models, any one of which can be taken as a null against which to compare more complex models: additive, multiplicative and stickbreaking ([57]). These models are similarly simple in that they all assume that fitness depends on the intrinsic effects of the constituent mutations. In the additive model, mutations have an absolute effect on the background fitness; in the multiplicative model their effect is proportional to the background fitness; in the stickbreaking model their effect is proportional to the distance between the background and the fitness optimum (generating diminishing returns). One advantage of these models is that because they lack higher order interactions ([58]) or phenotypic dimensions, they have few parameters to estimate. This benefit it not trivial because the amount of data available for model fitting is severely constrained: the full combinatoric network of *k* mutations contains 2^k^ − 1 observable effects.

We argue that modeling always benefits from the existence and use of meaningful null models. When null models are rejected in favor of a more complex one, the rejection is more than a straw man; rather, the way the complex model differs from the null(s) offers insights into the underlying biology. In other cases, we may find that the simple model provides a good enough approximation to be useful. The purpose of this work is to develop the methods for fitting and comparing the three basic landscape fitness models to data. Using simulations and empirical data, we then illustrate how to use these methods.

## 2. Methods

### 2.1. Overview

We begin by assuming that the data represent the complete set of 2*^k^* genotypes created from *k* mutations (wild type included). Later we return to the topic of other dataset structures. Our approach here is to fit the data to each of three models where the models have the same structure: the observed fitness (or phenotype) of each genotype is the expected fitness (or phenotype) under the specified model plus Gaussian error. The reader may notice that the Rough Mt Fuji model is analogous to the additive model used here ([35], [52]). Furthermore, the process of log-transforming data and then fitting it to the additive model (e.g. [52]) is analogous to assuming a multiplicative model (except a question remains about how to model the error; see below). Fitting the stickbreaking model in the same framework has not been done before. In order to do this, we must estimate the fitness boundary as well as the coefficients. After establishing how to fit the three models, we develop methods to compare them and identify the one(s) that best explain the data. We will use a Bayesian approach to assign posterior probabilities to the three models.

### 2.2. Notation

We begin by establishing some notation, much of which is standard. We will use capital letters to denote sets of mutations or random variables, it should be clear from the context whether we are referring to a set of mutation or a quantity which is observed with error (random variable). Let *K* = {1, 2, ⋯, *k*} be the set of all mutations under study. Let the set of mutations comprising a genotype be denoted by *G*, where *G* is a subset of *K* (*G* ⊂ *K*). We will use small letters to represent elements with in a set, or parameters of the model. For example we may refer to mutation *i* ∈ *G* as a single mutation among those in the set of mutations denoted by *G*.

### 2.3. Basic models

If there were no errors or noise in the model, then under the additive model, the fitness of genotype *G*, *w_G_*, would be
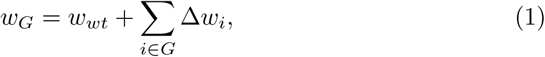

where Δ*w_i_* is the intrinsic effect of mutation *i* and *w_wt_* is the fitness of the wildtype. Fitness under the multiplicative model would be,
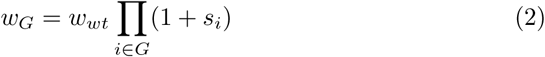

where *s_i_* is the intrinsic selection coefficient of mutation *i.*

In the stickbreaking model, the effect of a mutation is to close the distance to the fitness optimum by a proportion specified by its coefficient ([57]). Thus when a mutation has a stickbreaking coefficient of 0.25, it moves fitness 25% of the way to the fitness optimum. If the same mutation occurs on backgrounds of increasing fitness, the absolute effect of the mutation will diminish. Formally, the expected fitness under the stickbreaking model is given by
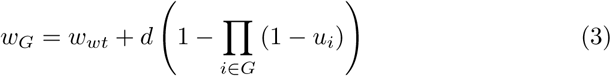

where *u_i_* is the intrinsic stickbreaking coefficient of mutation *i* and *d* is the fitness difference between the fitness boundary and the wildtype (see [57] for derivation of the stickbreaking model).

Even when one of these models is a valid, we expect real data deviate from the expected values for two reasons. First, the models are, at best, approximations of reality and deviations due to the underlying biological processes will exist. Second, there is experimental error in real data. We accommodate both of these sources of noise by combining them into one term such that the observed fitness of any genotype is given by its predicted fitness under the model plus a normally distributed error: *W_G_* = *w_G_* + *ϵ* where *ϵ* ~ N(0, *σ*^2^) and *w_G_* is given by equations 1, 2, and 3. We assume that the errors are independent across genotypes. Note that the stickbreaking and multiplicative models involve products instead of sums. This means they reside naturally on the log-scale while the additive model is on the non-log scale. It might seem appropriate, therefore, to model errors for stickbreaking and multiplicative as log-normal (normal on the log scale). Although we explored this possibility, we could not resolve the problem of how to compare models on different scales. A second reason to use a common error structure is that the experimental error will typically have a normal structure and this will not depend on the underlying model. Finally, using normal, and not log-normal, error allows us to use a maximum likelihood estimate of *d*.

### 2.4. Estimating distance to the fitness boundary, d, under stickbreaking

The first step in fitting the stickbreaking model to real data (which we do in the next subsection) is to estimate *d*. We develop three different methods of estimation.

#### Method 1: Maximum likelihood

Because we are assuming error is normally distributed, the maximum likelihood estimate (MLE) will be that value of *d* that minimizes the squared differences between observed fitness and the predicted fitness (right side of 3):
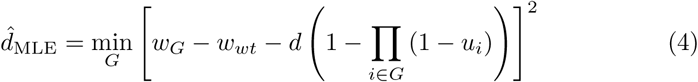

In practice, we find the MLE using the *optimize* function in R. All of the computational work in this paper is done the R environment ([59]).

#### Method 2: Relative Distance to Boundary (RDB) estimator

Equation(3) can be rewritten as
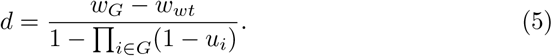

Notice that every genotype *G* the right-hand side of (5) gives a different way to represent *d*. The strategy is to begin by estimating Π_*i*∈*G*_(1 − *u_i_*) and then use that estimate in (5) to estimate *d*. The expression Π_*i*∈*G*_(1 − *u_i_*) represents the relative distance to the boundary (RDB) for genotype *G.* If genotype *G* produces a fitness gain of *w_G_* − *w_wt_* then distance to the boundary would be *d* − (*w_G_* − *w_wt_*) and the relative distance to the boundary would be 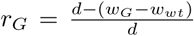. So it follows from (3) that
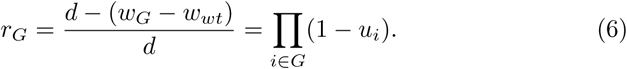

It would appear from equation (6) that in order to calculate the relative distance to the boundary *r_G_* using observable fitness effects, one would need to know *d* a priori. However, in the APPENDIX we obtain an expression for *r_G_* based on observable fitness effects independent of *d*. Equation (18) in the APPENDIX shows that

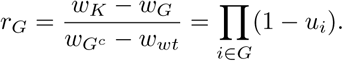

This leads to the following estimate 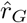 for the RDB of *G*
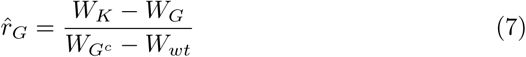

which leads to a set of estimates for the boundary *d* given by
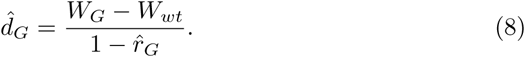

We now define the set of all estimates of *d* given by (8) to be
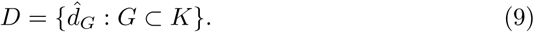

*D* can be viewed as a transformation of the fitness effect data. *D* contains 2*^k^* − 1 estimates of *d*. A measure of the center of the transformed data *D* will form our final estimate of *d*. Both the average and median of *D* produce valid estimates of the distance to the boundary *d*. We did extensive simulations on the properties of the mean versus the median and have concluded that for the noise levels explored here, the median estimator is the better alternative. Thus, this RDB estimator, which uses all genotypes, is
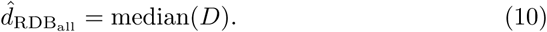

Notice that for data without error, *r_G_* can only fall between 0 and 1. With noise, however, genotypes can generate values outside this range. In particular, genotypes where *w_G_* < *w_wt_* or *w_G_* > *w_k_* cause problems. This led us to considered a modification to the RDB estimator where we use only the subset of *D* that come from genotypes where 0 < *r_G_* < 1, denoted *D*_01_. In the results section we will show that this modified estimator outperforms 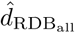. For notational simplicity we hereafter denote it as just 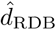. Formally then, the estimator is,
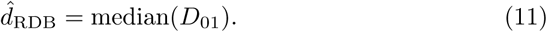

#### Method 3: Hybrid estimator

In order to do model selection (below), it is invaluable if we can fit the the stickbreaking model to every dataset, even if the fit is very poor. The two estimators just described do not always produce valid estimates of *d* and this prohibits fitting the stickbreaking model to every dataset. We define a valid estimate to be *d* > 0 and *d* < 10(*w_max_* – *w_wt_*) where *w*_max_ is the maximum observed fitness. An estimate < 0 implies a fitness boundary lower than the observed fitness values. The reason for not accepting values more than ten times the largest fitness difference from wild type (i.e. 10(*w*_max_ – *w_wt_*)), is because we want the stickbreaking model to be distinct from the additive model, yet the stickbreaking model approaches the additive model as *d* gets large and the coefficients get small ([57]).

What can be done when the MLE and RDB estimators both fail? One guaranteed way to always obtain an estimate of *d* is simply to use a value slightly larger (say 10%) than the largest observed fitness: 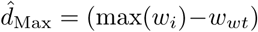 (1.1)) for all *i* ⊂ *G.* Note that defining *d* this way results in maximal coefficient estimates and generally a poor fit to the data. Our view is that if the data is so noisy and problematic that we cannot obtain a good estimate of *d*, then it is appropriate to disfavor the stickbrekaing model by using a small estimate of *d*. We suggest, then, the following rule as a way to estimate *d* across all datasets: use 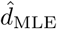 unless it fails to produce a meaningful estimate; use 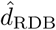 unless it fails; then use 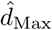. We justify this order in the results section below. We refer to this procedure for estimating *d* as 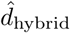.

### 2.5. Estimating coefficients

When mutation *i* is added to background *B* (where *i* is not in *B*), denote this genotype *B^i^*. Under the additive model, the expected value 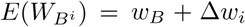, and hence a natural way to estimate the coefficient for mutation *i* is just 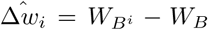 and average over all *B* and *i*. We make a small adjustment to this by weighting the observations on the wild type background twice as heavily as the other genotypes. This is because we assume wild type will generally serve as a control in fitness estimation with the consequence that it is observed more times and thus estimated much more precisely than the other genotypes. The consequence is that the variance of the difference 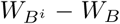 will be *σ*^2^ when *B* is wild type, and 2*σ*^2^ when *B* is not wild type. Weighting by the inverse of the variance, we get,
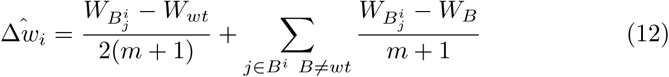

where *j* ∈ *B^i^* indicates sum over all genotypes containing *i*, *B* ≠ *wt* means exclude the case where *B* is wild type, and *m* is the number of backgrounds on which *i* appears (*m* = 2*^k^*^−1^ or half the genotypes in the dataset).

Because the multiplicative model involves a product, it is simplest on the log-scale. Taking the log of both sides of equation (2) and defining *y_G_* as the transformed fitness, we have
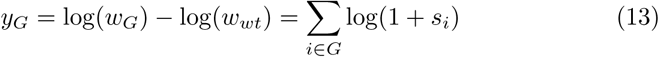

If we let *y_i_* = log(1 + *s_i_*), then equation 13 implies that 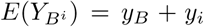. 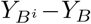 therefore provides an estimate of *y_i_.* We weight the estimates according to whether one or both backgrounds are observed with error and then transform back to the non-log scale:
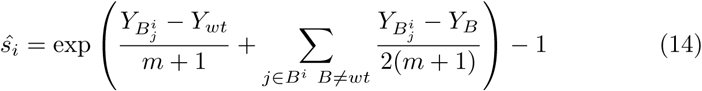

For stickbreaking, we also transform to the log-scale by taking the log of equation (3), rearranging, and defining *z_G_* as the transformed fitness,
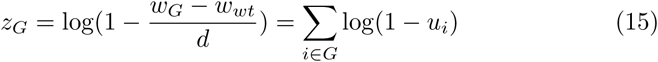

Letting *z_i_* = log(1 − *u_i_*) and replacing *d* with 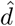 in equation (15), we see that 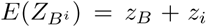 so that 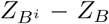 provides an estimate of *z_i_*. Again, we estimate *i* over all genotypes it appears in, weight by the inverse of the variance and then transform the estimate back to the non-log scale,
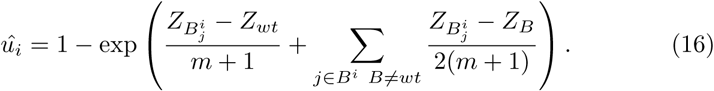

### 2.6. Estimating σ^2^

Recall that we assume all genotypes in the dataset except wild type depart from their predicted value as independent random normal deviates with mean 0 and variance *σ*^2^. We thus estimate *σ*^2^ by,
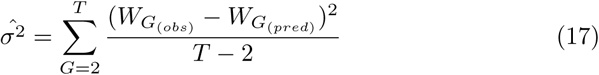

where 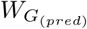 comes from substituting the estimated coefficients (and 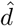 in the case of stickbreaking) into the appropriate equation (equation 1, 2 or 3).

### 2.7. Assessing fit and model selection

We are ultimately interested in determining which of the models (stickbreaking, multiplicative, or additive) are consistent with a set of data. Because it is straightforward to calculate the likelihood of the data under the three models, we first pursued using AIC to do model selection. To our surprise, this approach was unsuccessful. When we analyzed simulated data, we observed that under parametric conditions with low signal to noise ratios, the true model was falsely rejected an unacceptably high fraction of the time (i.e. ≥ 5%). We believe this owes to the nonstandard nature of the data as a network where each observation involves a different subset of parameters. We eventually abandoned AIC. Instead, we developed a Bayesian approach that creates a predictive model of posterior probability by training it on simulated data. The method has fours steps: (i) simulate data from priors, (ii) fit data to each model, estimate parameters and generate summary statistics, (iii) feed the summary statistics into a multinomial regression to train it, and (iv) use the multionomial regression model on other data (e.g. real data) to calculate the probability it comes from each of the three models.

We now cover these four steps (denoted i-iv) in greater detail. In step (i), we do simulations. We conducted separate simulations for networks with 3, 4, and 5 mutations. For each number of mutations, we simulate 10,000 datasets by drawing parameters from uniform prior distributions: each model (stickbreaking, multipliactive, additive) has equal (1/3) probability, a coefficient value (*u*, *s* or Δ*w* depending on the model) is sampled from a uniform (0.05, 0.5) and then assigned to all mutations in the dataset, and *σ* is sampled from a uniform (0.01, 0.1). For stickbreaking datasets, *d* = 1 throughout. We then simulate datasets according to the assumptions described in the ‘Basic Models’ section above.

In step (ii), we fit the data to each model. For stickbreaking, this requires first estimating *d*. For all three models, we then estimate the coefficients for each mutation. (Note, while we use the same coefficient value across mutations when simulating data, we estimate each one individually during the analysis.) We then summarize the fit using two statistics for each model: *R*^2^ and a P-score. *R*^2^ gives an overall measure of how close the predicted fitnesses are to the observed values under each model. It is obtained by calculating the mean fitness over the network (excluding wild type), taking the squared deviations from this mean, and summing to get the total sum of squares (TSS). We then estimate the coefficients (and *d* in the case of stickbreaking) and plug these estimates into equations 1, 2 and 3 to get predicted fitness values for every genotype in the network. We next take the differences between the predicted values and observed values, square them, and sum to get the residual sum of squares (RSS). Then, *R*^2^ = 1 − *RSS/TSS.* Note that this is not regression and there is no guarantee that the predicted values will be closer to the observed values on average than the overall mean is. Thus it is possible for poorly fitting models to generate *R*^2^ values < 0.

While *R*^2^ examines the nearness of observed and predicted values, the P-score assesses whether the pattern of deviations is consistent with a model. In short, if the data arose under the model being considered, then the observed fitness effects (as defined under that model) should not show a trend with increasing background fitness. If the data arose under a different model, they should should show a non-zero trend. To make this more precise, consider first the additive model. Let *B^i^* and *B* be a background with and without mutation *i.* By equation 1, 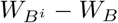 has expected value Δ*w_i_* regardless of what background is considered. Thus if the data arose under the additive model and we regress 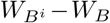 against *W_B_* for all *B*, we expect a line with intercept Δ*w_i_* and slope zero. If the data instead arose under the multiplicative or stickbreaking models, we expect positive and negative slopes respectively when we do this regression ([57]). The analogous argument for the multiplicative model leads to the conclusion that if the data arose under it, regressing 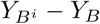 against *W_B_* should yields zero slope. If the data arose under the stickbreaking model, it is the regression of 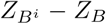 against *W_B_* that should show no slope. Note that this is the same approach pursued by [40], except that they only considered the additive model.

Our linear regression test to generate a P-score is therefore to take each mutation *i* = 1, 2, ⋯, *k*, consider each background upon which it appears, calculate the observed effect under each of the three models (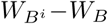 for additive, 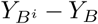 for multiplicative, and 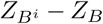 for stickbreaking), and regress these against *W_B_.* We then fit these data points to a simple linear model using least-squares and obtain a p-value. The information in the p-values (*p*_1_, *p*_2_, ⋯, *p_N_*) are then summarized by taking the sum of the logs of the p-values to yield a P-score: 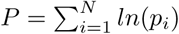. The smaller the p-values across mutations, the more negative the P-score becomes. Notice that the pattern of departure from zero under the incorrect model is is not actually linear ([57]). By assuming it is linear, we forego some power but benefit in terms of simplicity and computational speed.

Upon completing step (ii), the results are summarized as a matrix of 10,000 rows (one for each datset) by seven columns, one for the true model and six for the fit statistics: 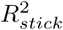, 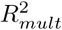, 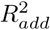, *P_stick_*, *P_mult_*, and *P_add_*. In step (iii), we use the matrix of results to do multinomial regression using the neural networks package in R, *nnet.* The multinomial regression uses six predictor variables (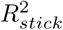,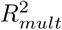, 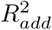, *P_stick_, P_mult_,* and *P_add_*) to to calculate the probability the dataset arose under each of the three models (stickbreaking, multiplicative, and additive). This is done separately for 3, 4, and 5. Once the model has been trained, it is ready to use on other datasets (step iv). To do so, a dataset is fit and summarized (step ii) and the summary statistics are passed into the previously trained multinomial regression model from step (iii) to yield posterior probabilities.

### 2.8. Incomplete networks

Not all datasets contain the entire network with all 2*^k^* genotypes formed from *k* mutations. One instance of this is simply when individual genotypes are missing from the network. Another case (which we refer to as a *double-mutant set)* is when the network contains just four genotypes: wild type, two individual mutations, and their combination double mutant. Suppose there are multiple such double-mutant sets. One possibility is that the mutations in each set are different (i.e. no mutations are shared). This generates an identifiabiltiy problem for stickbreaking (four datapoints yields three observed effects and there are three stickbreaking parameters to estimate) and we do not attempt to fit such data. Alterantively, it is possible that the same mutations appear across multiple double-mutant sets. In this case, we can view the data as a sample of the first and second steps of the much larger network. For example, in a bacteriophage dataset that we will analyze later, nine single mutations were engineered in various combinations to generate 18 double mutants. We think of this as 28 genotypes (including wild type) of the full 512 genotype network (2^9^ = 512).

Whenever the network is incomplete, a few minor adjustments to our approach are necessary. First, recall that the *RDB estimator* of *d* requires pairing a genotype with its complement (i.e. genotypes with and without a set of mutations). With incomplete sets, the necessary genotypes are often absent. When we cannot get RDB estimates of *d* for each mutation, we cannot employ this estimator. In this case, our approach is to use the MLE method if it exists, and largest observed fitness if it does not. Second, recall our method of model selection entails *P* scores that are based regressing effect size against background fitness. Regression requires three datapoints. If we do not have a mutation on three backgrounds, we cannot perform the regression. Thus sometimes one or more mutations will fail to yield p-values. Our approach is to base the *P* score on the mutations we do get p-values from. If we cannot get any p-values, we do model selection using *R*^2^ values alone. This approach is justified by the next adjustment. Third, we must rerun the model training simulations where we sample 10,000 datasets from our priors, but instead of using the full network as before, we use whatever data structure is observed in the real data. As before, we then fit each dataset to each model and then use multinomial regression to train a model for assigning posterior probabilities to the three models.

One word of caution about incomplete datasets is warranted. If a genotype is absent because it is inviable, then omitting it will bias the analysis. While one could assign such samples a fitness of zero, this will also introduce bias because, in reality, inviable genotypes represent a boundary condition that the models fail to incorporate.

## 3. Results & Discussion

The goal of this work is to compare the additive, multiplicative and stickbreaking models of epistasis. To do this, we first need to fit each of the three models to the data and second do model selection. For the additive and multipicative models, fitting is straightforward but for stickbreaking the distance parameter, *d*, must first be estimated. We therefore open with a subsection on estimating *d* and proceed to one on fitting data (i.e. estimating coefficients), then to model selection and finally to several subsections that deal with the analysis of different types of real data.

### 3.1. Estimation of d

To determine the best method of estimation, we simulated data under the stickbreaking model, setting *d* = 1, considering effect sizes that ranged from *u* = 0.1 to 0.5 in 0.1 increments, noise levels that ranged from *σ* = 0.02 to 0.1 in 0.02 increments, and complete genotype networks comprised of 3, 4, or 5 mutations. The results for a subset of illustrative cases with three or four mutations are presented in Figure 1. The MLE estimator fails most often, but when it does work, it is generally the least biased and has error at or below the others. Thus it is the estimator of first choice in the hybrid method. Between the two RDB (relative distance to boundary) estimators, the one that uses the subset of genotypes with an estimated boundary in the interval 0 to 1 (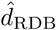; in black) outperforms the one that uses all genotypes (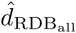; in grey). It fails far less frequently, tends to be less biased, and has similar rMSE; thus it is the second choice. The estimator based on the observed maximum fitness, 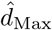, by definition never fails, but it is chronically biased low. It is the estimator of last resort. All estimators become good as signal to noise improves. The inset pie charts show that the hybrid estimator is dominated by the MLE and RDB estimators with the Max estimator only appearing when signal to noise ratio if very poor.

**Figure 1.**
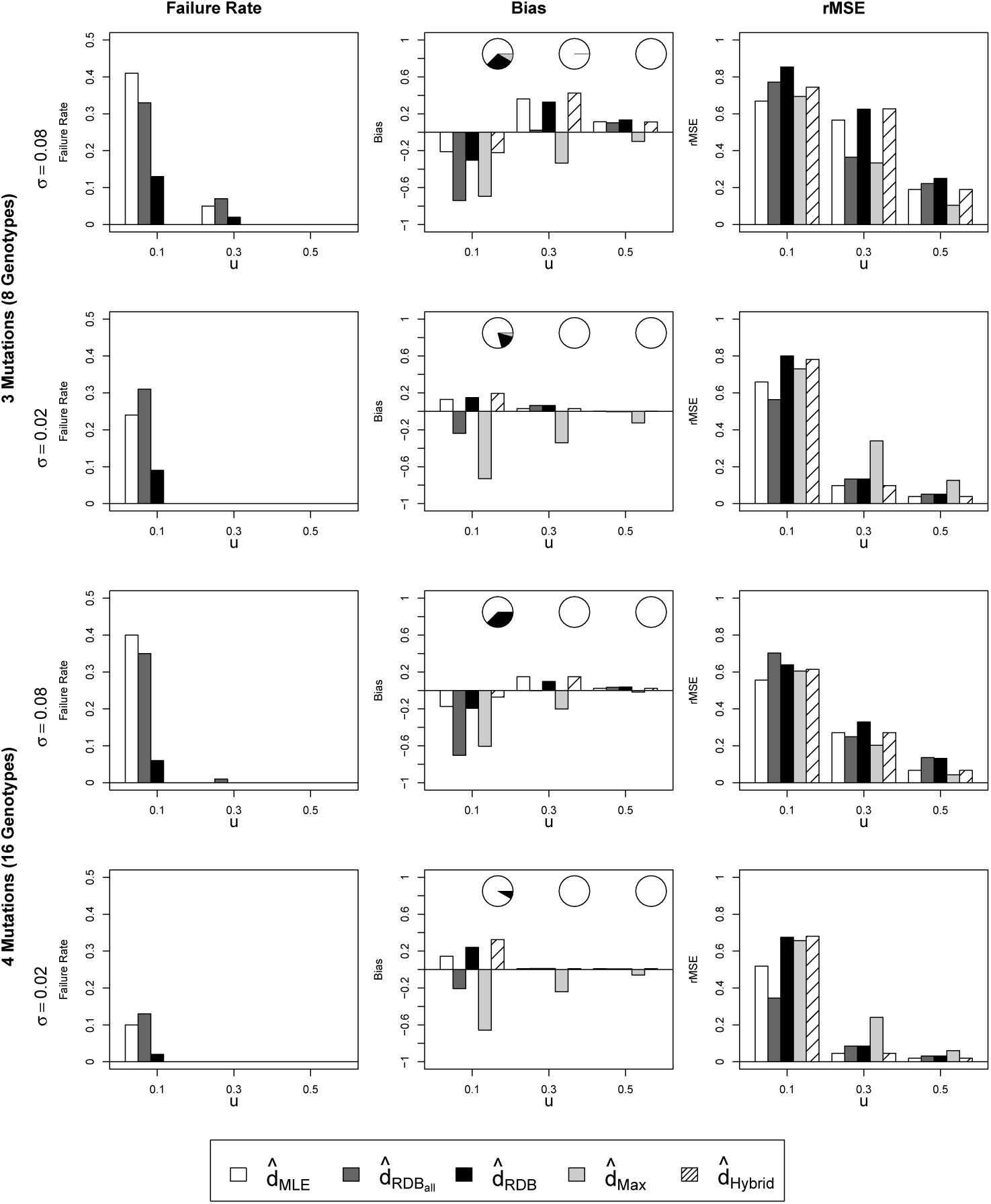
Failure rate (left column), bias (central) and root mean squared error (rMSE; right) for several estimators of the stickbreaking boundary, *d* (inset legend). Number of mutations and *σ* indicated to left of panels; coefficient size, *u,* on x-axis. Inset pie-charts show proportion of hybrid estimates based on MLE, RDB and Max estimators. Results based on 100 simulations per condition.

### 3.2. Coefficient estimation

Each of the three model has coefficients associated with each mutation that we estimate from the data. For stickbreaking, *d* is estimated first and then the coefficients are estimated based on 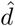. Figure 2 shows the 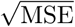 and bias for the stickbreaking coefficients based estimates of *d* from the hybrid method. The figure demonstrates three things. First, error and bias in estimates of *d* leads to substantial error and some bias in estimating *u.* Second, small effect sizes are associated with large proportional error: at *u* = 0.1 the 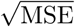 is also around 0.1. The errors as a proportion of the effect size are much smaller for = 0.3 and *u* = 0.5 where relative errors are more on the order of one-thrid and one-fifth respectively. Third, reducing noise (i.e. decreasing *σ*) has a greater effect than increasing the number of mutations. For example, when we hold *u* at 0.3 and compare 3 mutations and *σ* = 0.02 (small network, low noise) with 5 mutations and *σ* = 0.08 (large network, high noise) we see respective 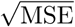 values of 0.037 and 0.063. Said another way, if there is a choice between generating a larger networks vs. reducing noise, reducing noise is the more effective way of getting good parameter estimates. Of course, if the noise is not experimental but biological, then it cannot be reduced. In comparing models, coefficient estimates under the additive and multipicative models have much smaller errors (Figure 3). Our simulations also confirmed that estimates under the multiplicative and additive models are unbiased (results not shown). This disparity between sickbreaking and the other models comes from the fact that the stickbreaking coefficients depend on estimating another parameter first while multiplicative and additive coefficients do not.

**Figure 2.**
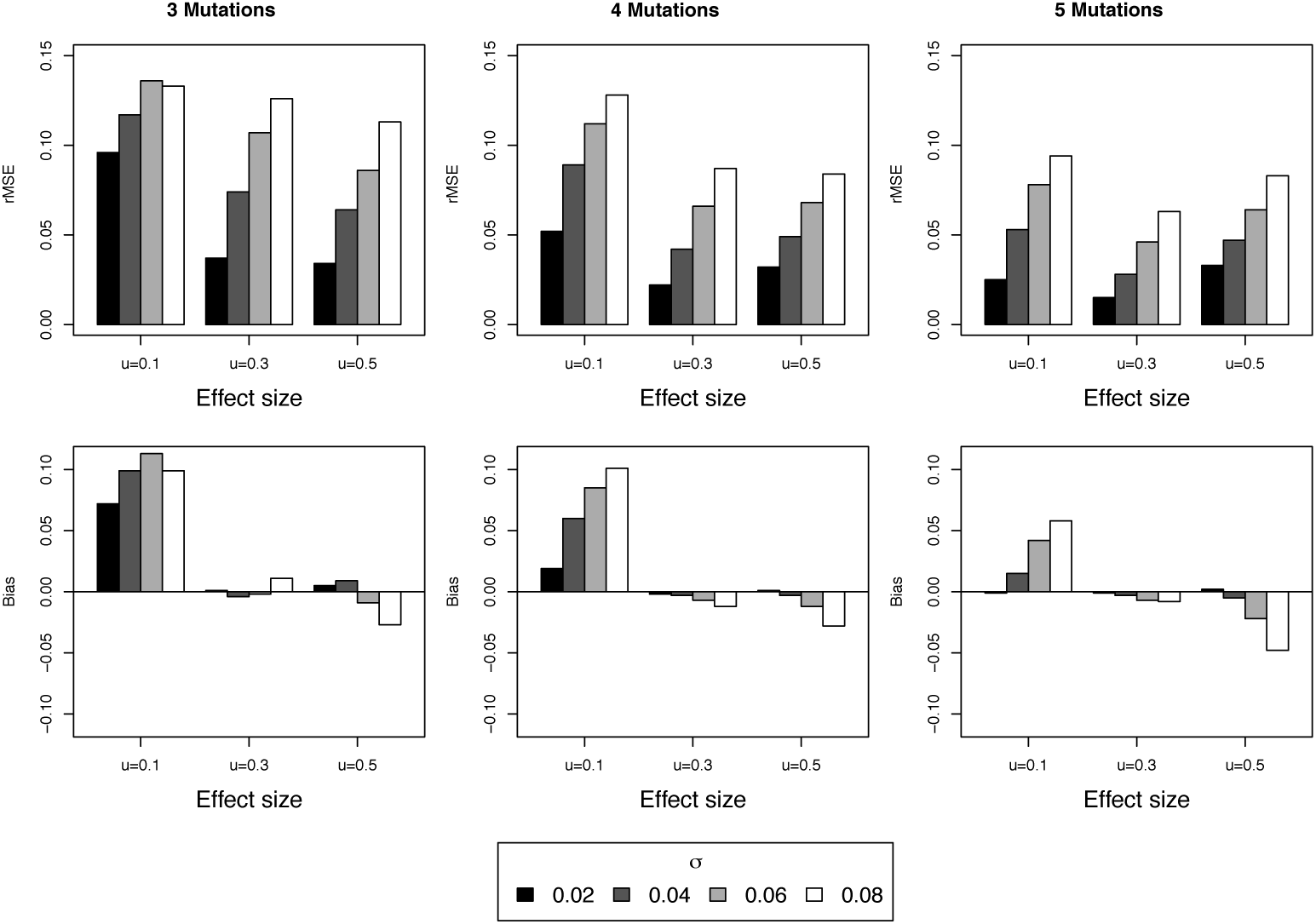
Root mean squared error (rMSE; top row of panels) and bias (bottom) of estimates of stickbreaking coefficients (*u*) as a function of number of mutations (columns of panels), effect size (x-axis) and *σ* (shaded bars, see legend at bottom). All estimates are based on the hybrid method of estimating *d.* Based on *d* = 1 and 1000 simulations per condition.

**Figure 3.**
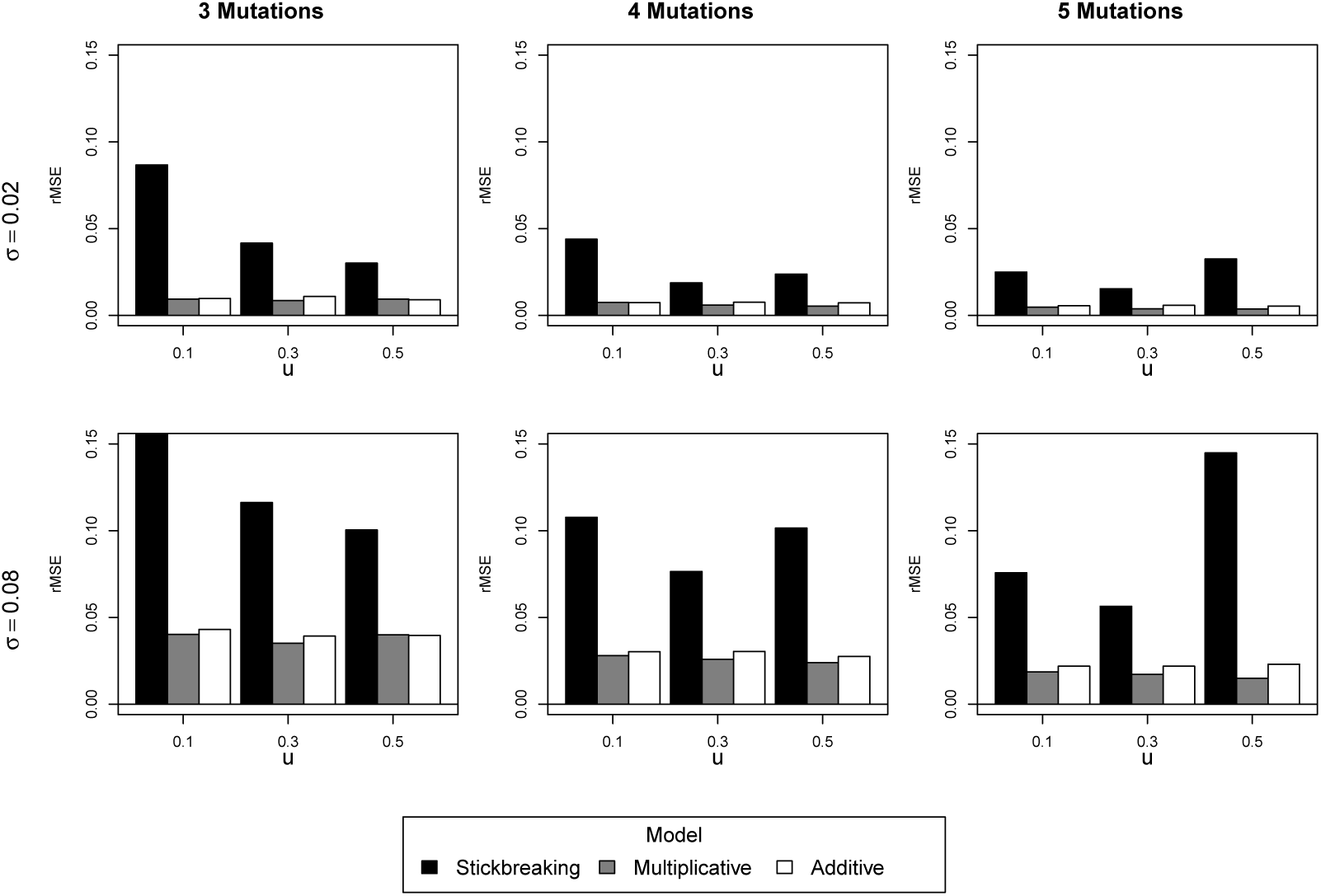
Root mean squared error (rMSE) of selection coefficients under all three models as a function of number of mutations (columns of panels), *σ* (top row 0.02, bottom 0.08), effect size (x-axis) and model (shaded bars, see legend at bottom). Based on 1000 simulations per condition.

### 3.3. Model selection

We are ultimately interested in identifying which of the three models best explains a given dataset. We took a Bayesian strategy for doing model selection in which we simulated a large number of datasets by sampling from prior distributions. Each dataset was then fit to each of the three models and summarized using *R*^2^ and a linear regression generated P-score (see Methods). We then passed these six measures of fit and the true model’s identity to a multinomial regression and allowed it to build a model that predicts the true model from the measures of fit. We did this for networks involving 3, 4 and 5 mutations seperately. The coefficients from this multinomial regression are presented in Table 1. We next generated test data. To do this we gridded parameter space: 3 true models (stickbreaking, multiplicative, additive) x 3 network sizes (3, 4, 5 mutations) x 5 coefficient values (0.1, 0.2, 0.3, 0.4, 0.5), and 4 *σ* values (0.02, 0.04, 0.06, 0.08). We then simulated 100 datasets per parameter combination. The performance of model selection was summarized as the mean posterior probability assigned to the true model and the proportion of replicates where the true model was rejected by having < 5% posterior probability.

**Table 1.** Coefficients for the multiple regression that produces posterior probabilities given six measures of fit: 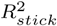, 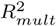, 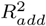, *P_stick_*, *P_mult_*, and *P_add_*.

| Mutations | Model | Intercept | $R^2_{stick}$ | $R^2_{mult}$ | $R^2_{add}$ | $P_{stick}$ | $P_{mult}$ | $P_{add}$ |
| --- | --- | --- | --- | --- | --- | --- | --- | --- |
| 3 | stick | -0.16 | 1.96 | -4.96 | 3.01 | 0.21 | 0.32 | -0.56 |
|  | mult | -0.05 | -0.08 | 12.51 | -12.61 | -0.03 | 0.54 | -0.60 |
| 4 | stick | -0.83 | 3.10 | -4.03 | 1.51 | 0.21 | 0.31 | -0.61 |
|  | mult | -1.09 | -0.24 | 18.84 | -17.32 | 0.00 | 0.45 | -0.57 |
| 5 | stick | -1.29 | 3.79 | -3.18 | 0.10 | 0.20 | 0.22 | -0.48 |
|  | mult | -1.54 | 0.65 | 26.48 | -24.54 | -0.019 | 0.36 | -0.43 |

This model selection method did a good job of limiting false rejections of the true model (type 1 errors). Of the 180 conditions tested, 175 of them had five or fewer false rejections in the 100 replicates. The remaining instances were scattered in parameters space and even here, error rates were not beyond what we would expect given a sample size of 100: two instances of 8 false rejections, two of 7 false rejections and one of 6 false rejections. The other critical part of model selection is how often it hones in on the true model and rejects others. Figure 4 shows the mean posterior probability of the true model under all 180 parameter conditions we studied. Surface regions in white highlight parameter space where the true model has posterior probability ≥ 95% while darker grey regions are those with lower posterior probability. White regions with high posterior probability correlate very closely with regions where the true model is uniquely identified a high frequency of the time. Two main trends jump out of these results. First, model selection is hard with only three mutation, better at four, and a lot better with five mutations. Stated more precisely, model selection leads to uniquely identifying the true model over a much greater range of parameter space as the number of mutations increases.

The second major pattern in the results is that the multiplicative model is the easiest to identify when true followed by stickbreaking and then additive. The multiplicative model ranks first because it produces the most distinct data: effect sizes for each beneficial mutation increase as background mutations accumulate even though (under our model) the error associated with them stays on the same scale. The stickbreaking model is opposite in that effect sizes shrink with accumulating mutations. While this leads to a distinct expected pattern in the data, two features of stickbreaking complicate things. One is the fact that the distance to the boundary, *d*, must be estimated from the data (unlike the other models). Two is the fact that while effects are shrinking with accumulated mutations, the error around them is not. The additive model ranks third simply because it produces patterns intermediate between the other two models. While data from the stickbreaking model is very rarely confused with the multipicative model and vice versa, data from the additive model can, by chance, resemble either stickbreaking or multiplicative.

**Figure 4.**
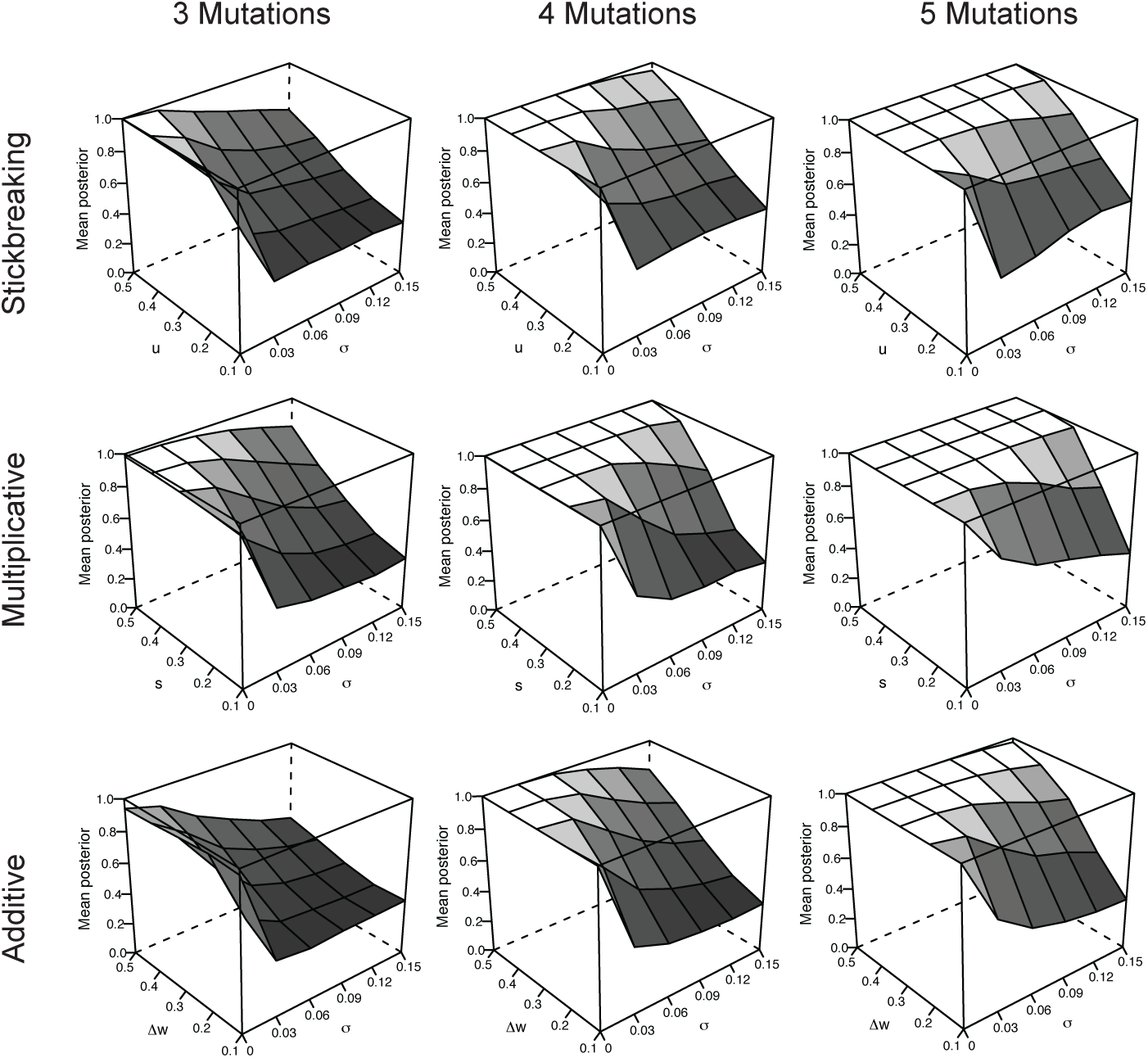
Mean posterior probability of the true model (z-axis in each plot) as a function of model (panel rows), number of mutations (panel columns), *σ* (x-axis) and evect size (*u*, *s* and Δ*w*; y-axis). Shaded white are regions with mean posterior probability ≥ 0.95.

### 3.4. Analysis of Real Data

To illustrate how our method may be implemented we selected several datasets from the literature. The first is from a study on fitness recovery in a *Methylobacterium* engineered with a foreign metabolic pathway that it must employ to grow on the sole carbon source of methanol. Nine mutations were identified over the course of adaptation. Four of these mutations were engineered in all combinations to form the complete 16 genotype network. The data fitted to each of the three models is shown in Figure 5A. When passed to the multinomial regression model, 99.1% of the posterior probability is assigned to the additive model. In their paper, Chou et al. developed an elegant cost-benefit model of the underlying metabolic processes, measured relevant phenotypes, and obtained a very good fit to the data. While their model provides more biological insight, the additive gives a very good approximation of the fitness effects observed among their mutations.

**Figure 5.**
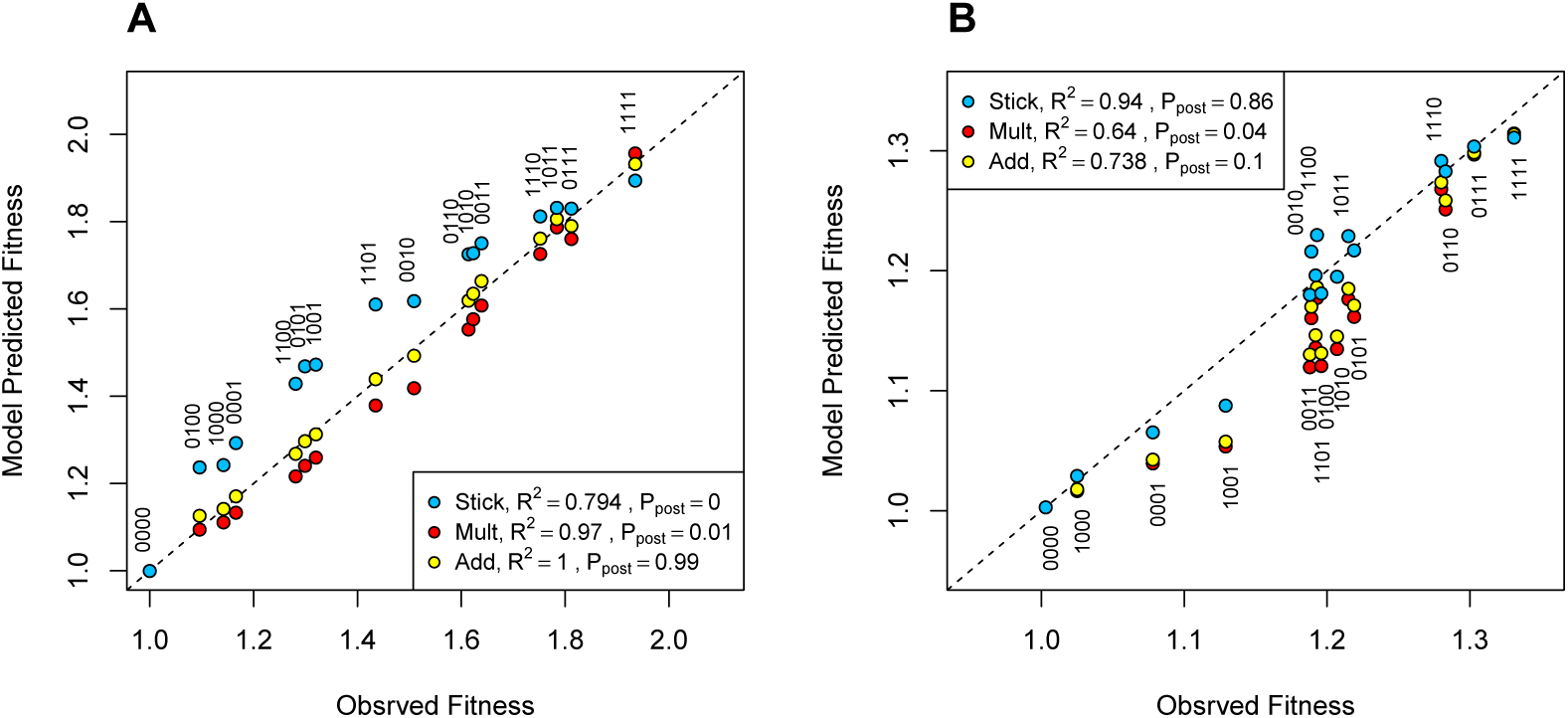
Observed vs model predicted fitness for two empirical datasets. In both cases, the inset legend indicates the model, *R*^2^ and posterior probability of model (*P_post_*). Each genotype in a dataset corresponds to a trio of vertically aligned circles (one per model) with the binary string indicating absence (0) and presence (1) of the individual mutations. (A) In the dataset from Chou et al. (2011) from *Methylobacterium,* the additive model fits very well and receives virtually all the posterior probability. The binary strings correspond to mutations *fghA*, *pntAB*, *gshA*, and *GB* in that order. In addition to the *R^2^* values shown in legend, the P-scores strongly favored the additive model: *P_stick_* = −24.7, *Pmult* = −20.3, and *P_add_* = −4.9. (B) In the Khan et al. (2011) data from *E coli,* the stickbreaking model fits best and receives 86% of the posterior probability, although the additive model cannot be rejected. As discussed in text, one outlier mutation (*pykF)* mutation was removed from the data before analysis. The binary strings correspond to mutations Δ*rbs*, *topA*, *spoT* and *glmUS* in that order. The P-scores also contribute support to the stickbreaking model: *P_stick_* = –0.59, *P_mult_* = –15.2, and *P_add_* = –12.2.

The second dataset we analyzed was an experiment by Kahn et al. (2011). Here the first five beneficial mutations in a long-term adaptation of *Escherichia coli* were engineered on the ancestral background in all 32 possible combinations. In their analysis, Kahn et al. examined additive fitness effects for each mutation as a function of background fitness. They showed that three of the five mutations in their dataset showed decreasing effects, one was not significantly different from zero, while one showed an increasing trend. These patterns correspond to our expectations under stickbreaking, additive, and the multiplicative models respectively. Not surprisingly, when we analyze the full 32 genotype network, we get ambiguous results with posterior probabilities for stickbreaking, multiplicative, and additive being 0.22, 0.40, and 0.38. We then removed the one strongly multiplicative mutation (+pykF) and reanalyze the 16 genotype network. When we do this we find that the data favors the strickbreaking model with 0.86 posterior probability compared to 0.10 for additive, and 0.04 for multiplicative. The fit of the data to the three models is illustrated in Figure 5B. Kahn et al. close their paper by stating that their results suggest a relatively simple epistasis function might be incorporated into models seeking to predict adaptation, though mutation +phkF demonstrates that there will be exceptions. Our results suggest that the stickbreaking model could provide exactly this type of simple function for approximating a common type of epistasis during adaptation.

### 3.5. Analysis of partial network data

Up to this point we have assumed the data covers all possible combinations of the studied mutations. However, this will not always be the case. In some instances there will be missing genotypes in the network. Another common type of dataset involves single mutants examined alone and in combination as the double mutant. If the same mutations are used across multiple double-mutant sets, then such data can be fit to the three models. A good example of this comes from work by Caudle et al. ([26]) on the bacteriophage ID11. Here, nine first-step beneficial mutations that arose during replicate adaptations at 37 C were engineered into 18 of the possible 72 double mutants; none of the higher order genotypes (eg. triples, quadruples, etc.) were created. Fitness was estimated at 33, 37 and 41 C. The datasets at 33 and 37 show such extensive sign epistasis that they do not fit any of the models considered here at all well (i.e. *R*^2^ values are <0). At 41C, however, sign epistasis was more moderate appearing in 7 of the 18 doubles, but being reciprocal (where both first steps are deleterious on the background of the other) in only two cases. When we fit the 41C data to the three model, we find that the stickbreaking model does a much better job than the others (Figure 6A). When the *R*^2^ and P-scores are passed to the multinomial regression model, 99.8% of the posterior probability is assigned to the stickbreaking model. This is not to argue that stickbreaking the best possible model here. Caudle et al. were able to achieve a considerably better fit to this data (*R*^2^ = 0.55 and 0.82) using more complex models that, in this cased, involved gamma-shaped phenotype-fitness functions. Nonetheless, this analysis illustrates that our approach can be used on this type of dataset that features only single and double mutants so long as sign-epistasis is rare.

**Figure 6.**
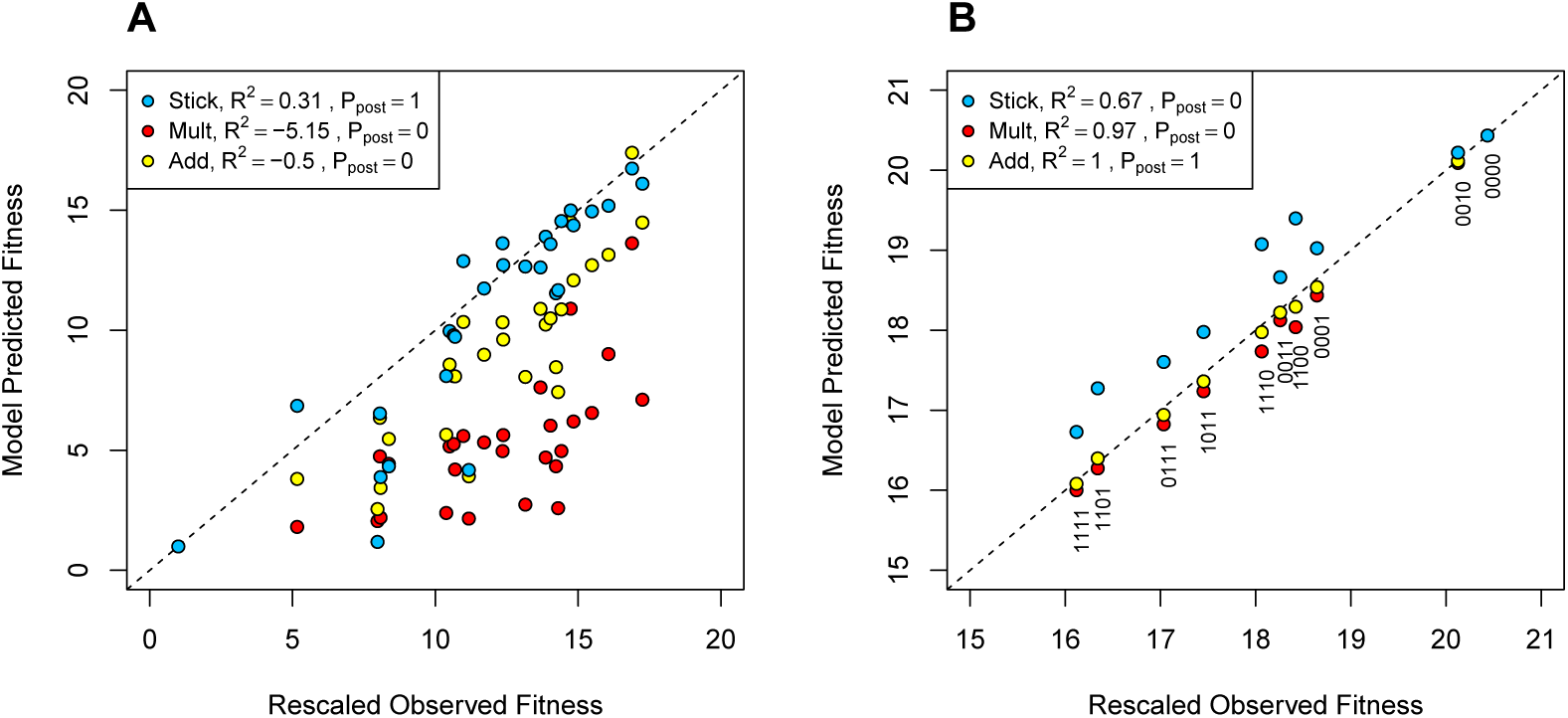
Observed vs model predicted fitness for subnetwork datasets. In both cases, the inset legend indicates the model, *R*^2^ and posterior probability of model (*P_post_*). (A) In the dataset from Caudle et al. ([26]) from the bacteriophage ID11, the stickbreaking model fits the data much better than the other two, although several of the single mutations to to the left have large errors. The P scores for the three models strongly contributed to the high posterior probabiltiy associated with stickbreaking: *P_stick_* = –11.5, *P_mult_* = −29.8, and *P_add_* = −29.9. (B) Combing synonymously recoded blocks of the poliovirus by Burns et al. ([60]) follows the additive model better than the other two. Each genotype in a dataset corresponds to a trio of vertically aligned circles (one per model) with the binary string indicating absence (0) and presence (1) of the individual mutations. The P scores also contributed to the additive posterior probability: *P_stick_* = −12.3, *P_mult_* = −3.3, and *P_add_* = −2.5

### 3.6. Analysis of deleterious data

Perhaps non-intuitively, our methods can also be used to analyze deleterious mutations, or even combinations of beneficial and deleterious mutations. We illustrate this by analyzing attenuation data from poliovirus. Burns et al. ([60])) recoded four contiguous capsid coding regions of 171-262 residues in length with synonymous mutations representing less preferred codons. They then created 10 of the possible 16 combinations of the recoded blocks and measured viral yield (plaque forming units or PFUs) over a 12 h growth period. We fit their data to the three models after log-transforming PFUs (since when growth is exponential, growth rate is proportional to the log of population size). The results, presented in Figure 6B, indicate that the additive model best describes the data, receiving over 99% of the posterior probability. This result is consistent with a result from the original paper where they found a strong, negative linear relationship between number of sites modified and PFUs.

## 4. Conclusion

We close with a few words about limitations and potential extensions of the framework advanced here. In terms of limitations, we have combined biological variance and experimental noise into a single variance term; in reality variance may differ among genotypes and there is generally information about how much of the noise is experimental (vs. biological) based on variance observed across replicates. This complexity could be added to our model in the future. Another limitation is that the interactions among all mutations are governed by the same model. Depending on the genes and mutations involved, this assumption may be violated (e.g. [40]). We experimented with developing a block version of our model motivated by Orr ([61]) where mutations are grouped into blocks of like-type and blocks interact. We ultimately ran into an overfitting problem, but if there were external information about how to group mutations or if networks were much larger than considered here, than strategy could be fruitful. A third limitation is that the model currently treats missing genotypes as simply absent. But if the genotypes are missing because they not not viable–something that will be especially common when mutations are deleterious– then the current approach is biased. To be done correctly, we need to treat inviable genotypes as having fitness censured by a lower boundary. This feature could be added to our model.

The methods and code presented here provide a framework for selecting among three basic landscape models. Sometimes, simple models are more use-ful than complex models when, for example, computational efficiency or mathematical simplicity are paramount. But the simplicity comes at a cost of course. These models cannot explain patterns like sign epistasis (except in treating it as noise) and when they do fit data well, they fail to provide a mechanistic explanation of it. We know that in reality mutations manifest their effects on fitness through their phenotypic effects. We are enthusiastic about modeling efforts that delve into phenotypic dimensions, including, for example, extensions of Fisher’s Geometric model (e.g. [6], [8]), models built on metabolic principles (e.g. [39], [46]), and models linked to protein stability (e.g. [62], [63]). We argue that the value and insight from these more complex models is far more compelling when the models they aim to improve upon are not straw men. We see one of the main extension of this work, therefore, as addressing how how the basic models fit here should be compared to more complex models. The tools provided here, we hope, make this and related uses of these basic landscape models readily accessible.

## Acknowledgements

This work was supported by the National Institutes of Health R01 GM076040, P20 GM104420, and P30 GM103324.

## Appendix

### Relative Distance to the Boundary (RDB)

Recall that *r_G_* defined by equation (6) has the following form for the stickbreaking model

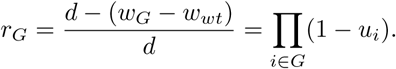

We obtain an expression for *r_G_* based on observable fitness effects independent of *d.* We do this by calculating the relative distance of genotype *G* from *K*, the genotype containing all available mutations. If we replace *d* with *w_K_* − *w_wt_* in the left hand side of the above equation and using equation (3) we get

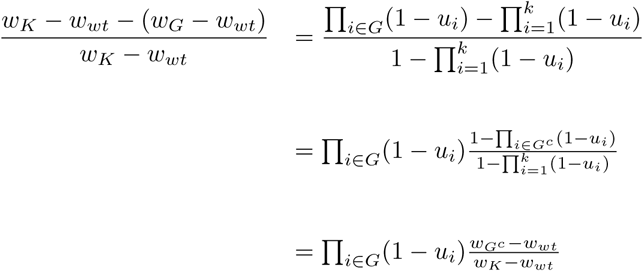

which implies
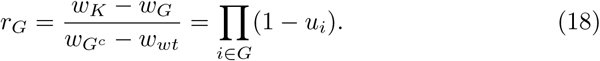

Equation (18) reveals that if one places genotype *G^c^* into the *G* background one obtains fitness effect *w_K_* − *w_G_*, which under the stickbreaking model is smaller than the fitness gain produced by placing *G^c^* into the wildtype background 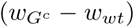. Comparison of the two fitness effects produces the RDB for *G.*

Note that equation (18) only applies for a proper subset *G* of *K* and cannot be used to calculate to calculate the RDB for *K.* However, we can still obtain an expression for *r_K_* by applying equation (18) to the genotype containing the single mutation *j* and the genotype containing all mutations but *j,* denoted by *j^c^*.
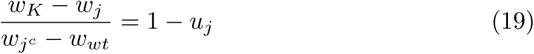

and
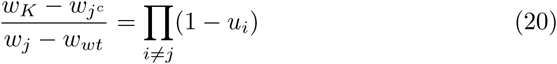

implying that
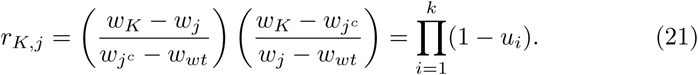

Note that *r_K_* = *r_K_*_,*j*_ for all *j*, but when we add noise to the mix, than the above will give us a set of estimates of *r_K_* for each *j*.

